# Carnivore-mediated seed crossings are more likely on roadkill hotspots

**DOI:** 10.1101/2025.10.01.679729

**Authors:** João Craveiro, Miguel N. Bugalho, Pedro G. Vaz

## Abstract

Roadkill hotspots concentrate animal movement and mortality, but whether they also interfere with animal-mediated seed dispersal—a conservation-relevant process underpinning plant regeneration and landscape connectivity—remains unknown. In Mediterranean oak woodlands of southern Portugal, we used seed mimics to test how road type (paved vs. unpaved) and road–forest context (edge vs. non-edge) shape carnivore-mediated seed crossings, and whether crossings coincide with roadkill hotspots detected by kernel density analysis. Bayesian logistic models indicated that seed-crossing probability was about twice as high on unpaved as on paved roads (predicted means 0.28 vs. 0.14), with weaker evidence for a negative edge effect. Crossing probability also tended to increase with carnivore abundance and distance to streams, and decrease with rodent density, albeit with some uncertainty. Crucially, paved sections intersecting roadkill hotspots showed nearly threefold higher predicted crossing probabilities than non-hotspot sections (0.51 vs. 0.18). These results yield clear conservation implications: prioritize paved segments that overlap carnivore roadkill hotspots. Implement precision measures—short-segment speed management, directional fencing that funnels animals to existing culverts/underpasses, and verge management that removes carcasses and limits small-mammal/scavenging attractants—to cut mortality while maintaining seed-mediated connectivity. A light, repeatable monitor–act–reassess workflow (hotspot mapping + seed-mimic trials + camera checks) enables rapid deployment and evaluation, and is transferable to other human-dominated forest mosaics. Hotspot-guided mitigation offers a pragmatic path to reconcile mobility infrastructure with living connectivity.

## INTRODUCTION

Mesocarnivores influence ecosystem functioning through diverse ecological interactions, including predation, scavenging, and seed dispersal through ingestion (Roemer et al. 2009; Herrera et al. 2015). Their capacity to disperse seeds stems from their large home ranges, broad movement patterns, and extended gut-retention times enabling them to transport seeds across long distances (Salgueiro et al. 2019). Yet the very traits that facilitate their broad dispersal simultaneously heighten their vulnerability to anthropogenic impacts (Craveiro et al. 2019; Azedo et al. 2022)—most notably roads, whose effects on wildlife are widely recognized (Benítez-López et al. 2010; Grilo et al. 2015; 2021; Ceia-Hasse et al. 2017; Quintana et al. 2022). Roads can reduce habitat connectivity by acting as physical and behavioural barriers, alter wildlife movement and habitat use, and cause direct mortality through vehicle collisions, with consequences for population persistence and gene flow (Forman and Alexander 1998; Coffin 2007; van der Ree et al. 2015; Quiles and Barrientos 2024). Yet how different roads shape carnivore-mediated seed crossings and whether high seed-crossing sections coincide with segments of elevated animal road mortality remains untested.

Roadkill hotspots arise from interactions among wildlife movement and habitat use, landscape context, and road and traffic characteristics (Bennett 2017; Pagany 2020; Dawson et al. 2025), and may also coincide with high-quality habitat (Clevenger et al. 2003; Grilo et al. 2011; Frangini et al. 2022). Beyond mortality, these segments may also undermine mutualistic interactions, including carnivore-mediated seed dispersal (Quiles and Barrientos 2024; Craveiro et al. 2025). Because these hotspots often overlap areas of regular animal use, they may also represent key seed-crossing points, affecting ecological connectivity (Cook and Blumstein 2013; Fabrizio et al. 2019). Thus, while hotspots are typically framed in terms of mortality risk, they may simultaneously function as risky corridors for seed flow—a dual role that remains unstudied and can have important implications for the conservation of both plants and animals.

Seed dispersal underpins plant recruitment and population dynamics (Nathan and Muller-Landau 2000; Wang and Smith 2002; Chen et al. 2019; Vaz et al. 2024). Carnivores that consume fleshy fruits transport viable seeds across landscape patches (Fedriani et al. 2023). In roaded environments, the interference of roads in seed dispersal is context dependent: both the road-forest context (edge vs. non-edge) and road type (paved vs. unpaved) shape movement and thus seed dispersal (Craveiro et al. 2025). Some carnivores avoid roads that delineate forest edges, reducing dispersal there, whereas others exploit edge habitats and move seeds into forest interiors (Tigas et al. 2002; Andersen et al. 2017; Craveiro et al. 2025). Unpaved roads—typically with lower traffic—are likely to show more carnivore-mediated seed crossings, whereas verges of paved roads can attract foraging via carcasses and prey, reinforcing road use (Ruiz-Capillas et al. 2013; Mata et al. 2017; Galantinho et al. 2020) and thereby enhancing dispersal in some contexts while elevating mortality.

Road mortality is the most visible consequence of road presence, often apparent as carcasses along roads (Santos et al. 2011). It imposes substantial population losses: vehicles kill hundreds of millions of vertebrates annually in the United States (Shilling et al. 2021) and ∼29 million mammals per year in Europe (Grilo et al. 2020). Carnivores are disproportionately affected—for example, 35% of annual Florida panther deaths are due to road collisions (Taylor et al. 2002); in Sweden, traffic kills 22000–33000 badgers and 6500–12500 red foxes each year (Seiler et al. 2004); and in southern Portugal, 47 medium-sized carnivores are recorded as roadkill per 100 km per year (Grilo et al. 2009). Because wildlife–vehicle collisions are often spatially aggregated along particular road segments (Santos et al. 2015; Costa et al. 2022; Pessanha et al. 2023), an open question is whether these mortality hotspots also represent areas of elevated carnivore-mediated seed crossings.

We conducted a field experiment in Mediterranean oak woodlands of southern Portugal to determine how road type (paved vs. unpaved) and road–forest context (edge vs. non-edge) shape the probability of carnivore-mediated seed crossings, and—on paved roads—to test whether road segments overlapping roadkill hotspots show higher seed-crossing probability. Specifically, we tested two hypotheses. H1: seed-crossing probability is higher on unpaved roads and in non-edge road-forest contexts, where traffic and edge-related openness are expected to impose a weaker movement barrier. H2: seed-crossing probability is higher on paved segments that overlap roadkill hotspots than on segments without hotspots.

## METHODS

### Study area and sampling design

We conducted the study on roads in the Alentejo region, southern Portugal (Fig. 1). The local climate is Mediterranean, with cold-wet winters and hot-dry summers (mean annual precipitation 620 mm; mean annual temperature 18 °C). The landscape consists of undulating terrain (200–400 m a.s.l.) dominated by *montado* (also denominated as *dehesa* in Spain) and pastures. *Montado* is characterized by open-canopy cork (*Quercus suber*) and holm (*Q. rotundifolia*) oak stands with an understory of shrubs (e.g., *Cistus*, *Ulex*, *Erica*, *Lavandula* spp.; Bugalho et al. 2009, 2011; Castro and Freitas, 2009; Azedo et al. 2022) and grassland areas. In Mediterranean systems such as evergreen oak woodlands of the Iberian Peninsula, where oak decline and limited natural regeneration have been documented (Bugalho et al. 2011; Vaz et al. 2019; Wadud et al. 2024), carnivore-mediated seed dispersal is particularly relevant (Herrera et al. 2015; Craveiro et al. 2025).

**Fig. 1.**
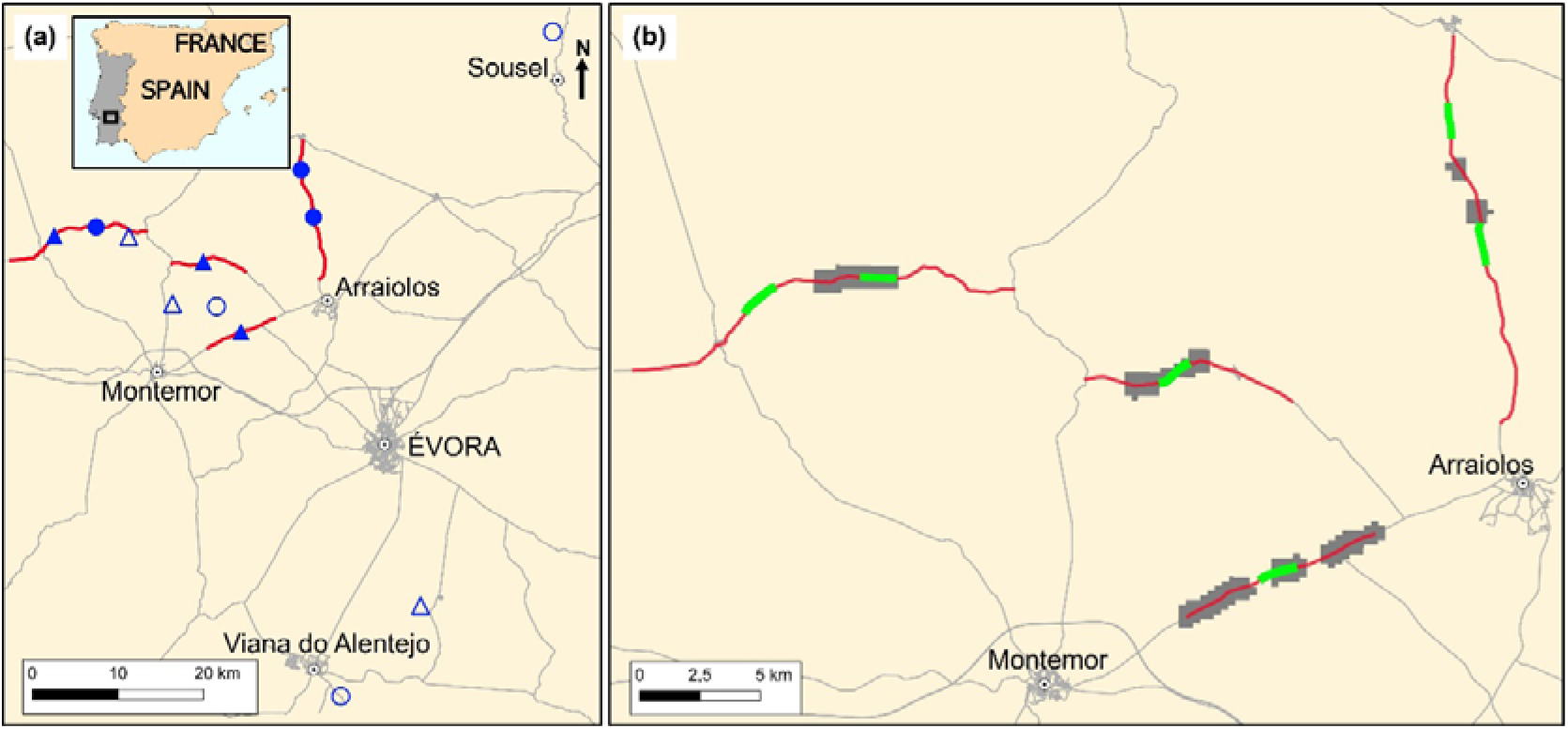
Study area and sampling layout in the Alentejo, southern Portugal. (a) Regional context and locations of the twelve 1.2-km road sections where carnivore-mediated seed crossings were monitored. Symbols encode road–forest context (circles = edge; triangles = non-edge) and road type (filled = paved; open = unpaved). Red lines indicate the National-Road transects driven weekly for roadkill surveys, which encompassed the six paved 1.2-km sections. (b) Detail of the six paved 1.2-km sections (light-green lines) and carnivore roadkill hotspots (grey patches; KDE, 500-m bandwidth), showing that three sections intersected one hotspot each and three intersected none.

We selected 12 road sections (1.2 km each; 3.1–32.8 km apart; mean = 8.7 km) encompassing two road-forest contexts—six bordering oak forests (edge) and six traversing oak forests (non-edge)—and two road types (six paved, six unpaved) (Fig. 2). Edge roads were characterized by oak woodland on one side and open pasture or cropland on the other. Paved roads (∼8 m wide; range: 6–11 m) were National Roads with asphalt pavement and a traffic volume of 95 vehicles/hour (range: 19–234) corresponding to 71 vehicles h□¹ (95% CI 16–125) on edge roads and 99 vehicles h□¹ (95% CI 19–392) on non-edge roads (see www.infraestruturasdeportugal.pt). Unpaved roads (∼4 m wide; range: 3–7 m) were public dirt roads with low traffic, mainly for agricultural access. To account for spatial variation, we established three replicates per factor (2 road-forest contexts × 2 road types × 3 replicates = 12 road sections; Fig. 1a).

**Fig. 2.**
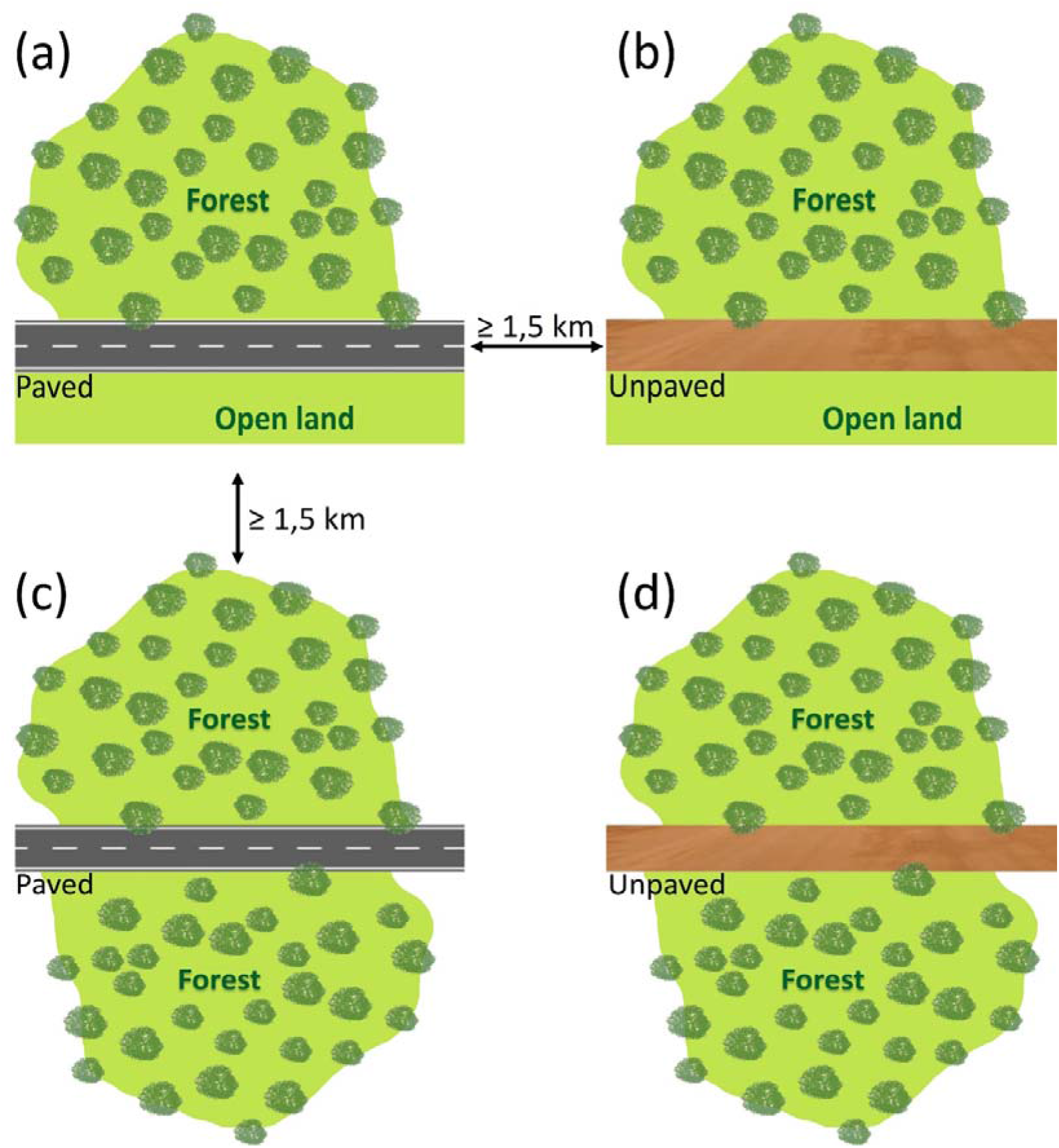
Schematic representation of the road-forest contexts and road types. (a, b) Edge sites with different road types, where the forest borders a tree-free open area, typically used for pasture or cropland. (c, d) Non-edge sites with different road types traversing a continuous forest.

Several carnivore species in the study area contribute to fleshy-fruit dispersal, including red foxes (*Vulpes vulpes*), European badgers (*Meles meles*), and stone martens (*Martes foina*), all of them moving seeds of brambles (*Rubus ulmifolius*), common hawthorns (*Crataegus monogyna*), and Phoenician juniper (*Juniperus phoenicea*) (Rosalino et al. 2010; Peredo et al. 2013; Farris et al. 2017). Other seed-dispersers include Egyptian mongoose (*Herpestes ichneumon*), Common genet (*Genetta genetta*), and Eurasian otter (*Lutra lutra*) (Craveiro et al. 2019).

### Seed-crossing sampling and covariates

To assess the influence of road-forest context, road type and covariates on seed crossing probability, we supplied dried common figs (*Ficus carica*), which have been proven suitable in feeding experiments (González-Varo et al. 2013; Herrera et al. 2015; Craveiro et al. 2025). Figs were offered once (no replenishment) across the 12 road sections from October 4, 2022 (fall) to July 18, 2023 (summer), ensuring proportional representation of each road-forest context and road type to avoid seasonal biases (Craveiro et al. 2025).

Within each road section, we installed six supply stations on each side of the road, approximately 200 m apart; each station was positioned 5 m beyond the verge, totaling 144 stations. At each station, we placed six aluminum trays (14.5 × 12.0 × 4.0 cm) spaced 10 m apart, each containing six dried figs. Every fig embedded 10 plastic beads uniquely color-coded for each station (seed mimics, 2.6 mm in diameter) (Herrera et al. 2015; Salgueiro et al. 2019). To document carnivore consumption of figs, we placed two Bushnell CORE S-4K model 119949C cameras at two randomly selected stations.

Eight days after deployment, we systematically searched for carnivore scats containing seed mimics within a 1200 m radius buffer around the supply stations, corresponding to the average home range radius of the carnivore community (Dekker et al. 2001; Planillo et al. 2018; Craveiro et al. 2019, 2025); this timing followed pilot trials indicating that recoveries within buffers stabilized by ∼8 days. The search focused on carnivore-utilized paths (e.g., trails, firebreaks, animal tracks; Herrera et al. 2015), covering on average 88% of the paths per buffer (range: 81%–98%). To reduce bias in detectability, all paths were traveled by the same observer walking at a constant mean speed of 2 km h□¹. Scat locations were georeferenced using a Garmin Dakota 10 GPS (mean error ± 3 m). Seed mimics from each scat were matched to their station of origin via the color code, allowing us to determine whether a ‘seed’ crossing occurred (scat on the opposite side of the road from its supply station; no scats were found on the road) and to calculate the perpendicular distance to the road verge. Although scat detection is imperfect, our standardized, path-based protocol and identical effort across sections could reduce heterogeneity in detectability and thus minimize bias in among-section comparisons of carnivore-mediated seed-crossing probability.

Tree density (oak trees/ha) was estimated within each 1200-m buffer around the supply stations, as this may influence carnivore habitat use (Cagnacci et al. 2004; Craveiro et al. 2025). Using GIS, we generated five random points per road side and created 50-m radius buffers around—twice the standard plot radius employed in Portugal’s National Forest Inventory, which improves tree detection on satellite imagery—, totaling 10 buffers per site. Oak density was calculated by dividing the satellite-derived count of oaks in the buffers by their combined area, obviating exhaustive counts across the 1200-m buffer.

Carnivore scats were assigned to species, where possible, using a combination of morphological and contextual characteristics, including overall shape, extremity form, texture, segmentation, color, odor, and deposition location (e.g. latrines, elevated sites, trails, or conspicuous marking locations; Brown et al. 2003; Olsen 2013). Because species assignments were not genetically confirmed, they may be subject to identification uncertainty and were used only for descriptive summaries; all carnivore species were pooled in the seed-crossing analyses. To account for variations in carnivore activity within the buffer, we calculated a Kilometer Abundance Index (KAI) as scats km□¹ along walked paths (Vincent et al. 1991), as a proxy for carnivore abundance. Additionally, we quantified landscape characteristics within the 1200-m buffer using QGIS (v3.20.3-Odense), considering total stream length, distance to nearest stream, the number and total area of water patches—variables selected a priori for their ecological relevance to carnivore-mediated seed dispersal and because they could be quantified consistently for all sections.

To assess rodent effects on carnivore-mediated seed crossings, we conducted a 4-night live-trapping survey at 12 rodent sampling points per 1.2-km road section. Six points were placed along the road (three on each side, 3–10 m beyond the outer edge of the verge), and six points were placed in the forest interior on one side of the road, arranged along a linear interior transect to minimize road influence. All sampling points were spaced >200 m apart. At each point, we deployed four Sherman traps along a short transect perpendicular to the road (or to the interior transect) at 0 m, 10–15 m, 20–30 m, and 40–50 m from the point center, placing traps in favorable microhabitats to maximize captures (under *Cistus* or *Ulex* shrubs or in tall grass; Vaz et al. 2024). Traps were baited with peanut butter on toast (4 × 4 cm), provided with cotton wool for insulation, checked at sunrise each day, and left in the same locations throughout the session. Captured rodents were marked with a small lateral fur clip (Hoffmann et al. 2010; Jumeau et al. 2017) and released at the capture site. Trap effort was 192 trap-nights per road section (12 points × 4 traps × 4 nights; total effort 2304 trap-nights). Procedures complied with Directive 2010/63/EU on animal use in research.

Rodent density by road section was estimated with Rcapture (v1.4-4; Baillargeon and Rivest 2009) under a closed-population assumption (no mortality or movement during the 4-night session), using the bias-corrected closedp.bc estimator for capture–mark–recapture data. Individual heterogeneity in capture probability and behavioural responses to trapping (e.g. trap-shyness or trap-happiness) were not explicitly modelled. Rodent density estimates were therefore interpreted as measures of relative rodent availability for comparisons among road sections and may contain uncertainty arising from unmodelled variation in capture probability.

### Roadkill surveys and hotspot detection

Between August 11, 2022, and August 10, 2023, we conducted weekly roadkill surveys (every Thursday) of carnivore species along four road sections (8.5–17 km in length; mean 12.8 km; Fig. 1b) (Grilo et al. 2009; Santos et al. 2015). To evaluate the spatial relationship between roadkill hotspots and seed-crossing sections, each survey section included the previously defined 1.2-km paved segments, yielding 51 km of roads surveyed weekly (total survey effort = 2652 km; 51 km × 52 visits). A single observer conducted surveys at sunrise, driving at ∼50 km·h□¹ (following standard roadkill survey protocols; e.g., Santos et al. 2018) and stopping to georeference each carnivore roadkill with a Garmin Dakota 10 GPS (mean positional error ± 3 m). Carcasses were removed after recording to avoid double counting.

We detected roadkill hotspots using kernel density estimation (KDE) (Bíl et al. 2013; Medinas et al. 2021) with a fixed 500-m bandwidth, previously recommended for identifying roadkill clusters and planning mitigation at the road scale (Grilo et al. 2009; Carvalho and Mira 2011; Morelle et al. 2013; Santos et al. 2016). Statistical significance was evaluated against complete spatial randomness via Monte Carlo simulations (n = 999; α = 0.05; Bíl et al. 2013). Given the small number of hotspots and their occurrence on separate road sections, network-wide autocorrelation tests (e.g., Moran’s I) were not informative (Carrijo and da Silva 2017). For subsequent analyses, we treated hotspots as operationally independent, because they occurred on different roads and were >500 m apart (i.e., farther than the KDE bandwidth). Hotspots were identified with the sparr package in R (Davies et al. 2017). We overlaid the KDE significance map on the six 1.2-km paved sections, classifying each section as hotspot-present or hotspot-absent (3 present, 3 absent). This binary factor was then used as a predictor in the model testing hotspot-presence effect on seed-crossing probability (see §2.4).

### Analyses of seed-crossing probabilities

To address both questions (effects of road attributes and of hotspot presence on seed-crossing probability), we modelled whether a seed crossed the road (1/0) with a Bernoulli model and logit link (Bayesian logistic regressions fit in brms (Stan; Bürkner 2017).

For the analysis of road attributes vs. seed-crossing probability, road type (paved vs. unpaved) and road–forest context (edge vs. non-edge) were specified a priori given our primary hypotheses, and their interaction was evaluated. We also considered a section-level random intercept (1|site; “site” denotes each 1.2-km road section) to account for within-section clustering. Additional covariates were distance to road, carnivore abundance (KAI), total stream length, distance to nearest stream, rodent abundance, number of water patches, area of water patches, and tree density (Table 1). Numeric predictors were centred and scaled. For a parsimonious model, we compared the interaction model with an additive specification (no interaction) and then tested removing each covariate one at a time (road type and road–forest context were retained a priori throughout). We interpreted leave-one-out differences in expected log predictive density (ΔELPD) whose absolute value exceeded 2×SE as meaningful (e.g., Vaz et al. 2021). This supported a final additive model (road type + road-forest context, no interaction) that omitted total stream length, number of water patches, and area of water patches (Table 3). Variance inflation factors (VIF; adjusted GVIF for categorical predictors) were < 2.36 overall and < 1.70 in the final model, indicating low multicollinearity (Lüdecke et al. 2021). For the analysis of hotspot presence vs. seed-crossing probability, carnivore roadkill hotspot presence (yes/no; §2.3) was the sole fixed predictor, and we considered a section-level random intercept (1|site) as above.

**Table 1.** Names, definitions, and range of variables considered in the analysis of carnivore-mediated seed crossings.

| Variable | Description | Range |
| --- | --- | --- |
| Road-forest context | Position of the road relative to the forest: "edge" (oak forest on one side, open area on the other) or "non-edge" (road traversing oak forest) |  |
| Road type | Road surface type: "paved" (asphalt) or "unpaved" (dirt road) |  |
| Distance to road | Shortest distance (m) from the seed's final location within a carnivore scat to the road. |  |
| Carnivore abundance | Kilometric abundance of carnivore scats | 0.69–3.01 |
| Rodent density | Estimated (individuals/ha) within each site | 1.91–7.14 |
| Length of streams | Total length of watercourses (m) within a 1200-m buffer. | 4103–9597 |
| Distance to stream | Shortest distance (m) from the seed's final location within a carnivore scat to a stream. | 45.91–1209.65 |
| Number of water patches | Number of water patches. | 12–32 |
| Area of water patches | Total area (ha) occupied by water patches within. | 1.68–20.74 |
| Tree density | Number of oaks per hectare estimated from ten 50-m radius buffers (five per road side). | 17.60–55.40 |

For both models, to aid convergence and limit overfitting, we assigned weakly informative priors to all effects (e.g., Gelman 2025). For the road-attributes model we used Normal(0, 10) priors for all β coefficients except road–forest context, for which we used Normal(0, 13); for the hotspot model we used Normal(0, 13) for the hotspot presence coefficient. We ran MCMC with 4 chains × 4000 iterations (1000 warm-up), yielding 12000 post-warm-up draws. We checked convergence using trace plots and RO ≤ 1.01, with adequate bulk/tail effective sample sizes; posterior predictive checks indicated good fit. Analyses were performed in R v4.5.0 (R Core Team 2025). For Bayesian statistical inference, we reported model coefficients (*β*) parameters of the models, their credible intervals, and tested statistical hypotheses (e.g., McElreath 2018). We report directional posterior probabilities [Pr(*β* > 0) or Pr(*β* < 0)] as an evidence measure. We describe values ≥ 0.95 as strong, 0.80–0.95 as moderate, and < 0.80 as weak evidence for an effect, while avoiding dichotomous ‘significance’ thresholds (Hobbs and Hooten 2015; Kruschke 2015; McElreath 2018).

## RESULTS

Across approximately 350 km of systematic scat searches along the 12 sections, we recorded 612 carnivore scats; 41 contained seed mimics, yielding 59 dispersal events—some scats contained seed mimics from multiple stations—, 27% of which were seed road crossings (Table 2). Crossings were more frequent on unpaved vs. paved roads (11 vs. 5; 2.2×) and in non-edge vs. edge road–forest contexts (11 vs. 5; 2.2×). Dispersal events were attributed to stone martens (36%), European badgers (34%), and red foxes (30%). Among crossings, red foxes mediated 44%, followed by stone martens (31%) and European badgers (25%). Camera traps logged 359 animal detections, 44% of which were carnivore detections. Among these, red foxes accounted for 49%, stone martens 39%, common genets 9%, Egyptian mongooses 2%, and European badgers 1%. Of the 153 fig removals, ∼70% were attributed to carnivores; among these, 52% were red foxes, 45% by stone martens, and 3% by Egyptian mongooses.

**Table 2.** Counts of carnivore-mediated seed dispersal events (“dispersals”) and of events that crossed the road (“crossings”), by road–forest context (edge vs. non-edge) and road type (paved vs. unpaved), and—for paved sections only—by roadkill hotspot presence (absent vs. present). Totals refer to the 12 sections (six edge, six non-edge; six paved, six unpaved). For hotspot presence on paved roads, three sections were classified present and three absent. “Crossing” = scat with seed mimics deposited on the side of the road opposite its supply station.

| Road-forest context |  | Road type |  | Total |
| --- | --- | --- | --- | --- |
|  |  | Paved | Unpaved |  |
| edge | dispersals | 4 | 13 | 17 |
|  | crossings | 2 | 3 | 5 |
| non-edge | dispersals | 13 | 29 | 42 |
|  | crossings | 3 | 8 | 11 |
| *Total* | dispersals | 17 | 42 | 59 |
|  | crossings | 5 | 11 | 16 |
| Paved sections: roadkill hotspot presence |  |  |  |  |
| absent | dispersals | 11 |  |  |
|  | crossings | 2 |  |  |
| present | dispersals | 6 |  |  |
|  | crossings | 3 |  |  |

Among our paved 1.2-km sections, the proportion of dispersal events that crossed the road was 2.7× higher in sections with roadkill hotspot presence (50%; see §3.2 for hotspot detection details) than in non-hotspot sections (18%) (Table 2). During weekly surveys we recorded 33 carnivore roadkills—65 per 100 km yr□¹ (Table A.1, Appendix A)—of which 88% were adults. Fifteen occurred within KDE-defined hotspots intersecting the three paved 1.2-km sections where seed crossings were monitored. Within these hotspots, badgers were most affected (6, 40%), followed by genets (3, 20%) and, with two deaths each (13%), red foxes, Egyptian mongooses, and stone martens. No carnivore roadkills were detected on the three non-hotspot paved seed-crossing sections. Outside the six paved 1.2-km seed-crossing sections—that is, elsewhere along the weekly roadkill transects—red foxes were most frequently killed (9 of 18; 50%).

During the live-trapping survey, we captured 57 unique rodents, 28 of which were recaptured at least once. Captures comprised Wood mouse, *Apodemus sylvaticus* (80%), and Algerian mouse, *Mus spretus* (20%), across the 144 sampling points. Closed-population capture–recapture estimates (Rcapture) yielded a mean density of 3.72 rodents ha□¹ (range 1.91–7.14 ha□¹) across the 12 sections; we used these estimates for among-section comparisons of relative rodent availability.

### Carnivore-mediated seed crossing probability

Accounting for nesting with a section-level random intercept, our Bayesian mixed-effects logistic model (Table 3; Fig. 3) indicated that road type influences the probability of carnivore-mediated seed crossings. Consistent with our hypothesis, directional Bayesian tests for the road-type contrast supported higher crossing probabilities on unpaved than on paved sections, with an estimated 85% posterior probability for this direction of effect. Predicted mean crossing probabilities were 0.28 (unpaved) vs. 0.14 (paved), doubling on unpaved sections. In contrast, evidence for a road–forest context effect was limited, with an estimated 65% posterior probability that crossings were higher in non-edge than edge contexts.

**Fig. 3.**
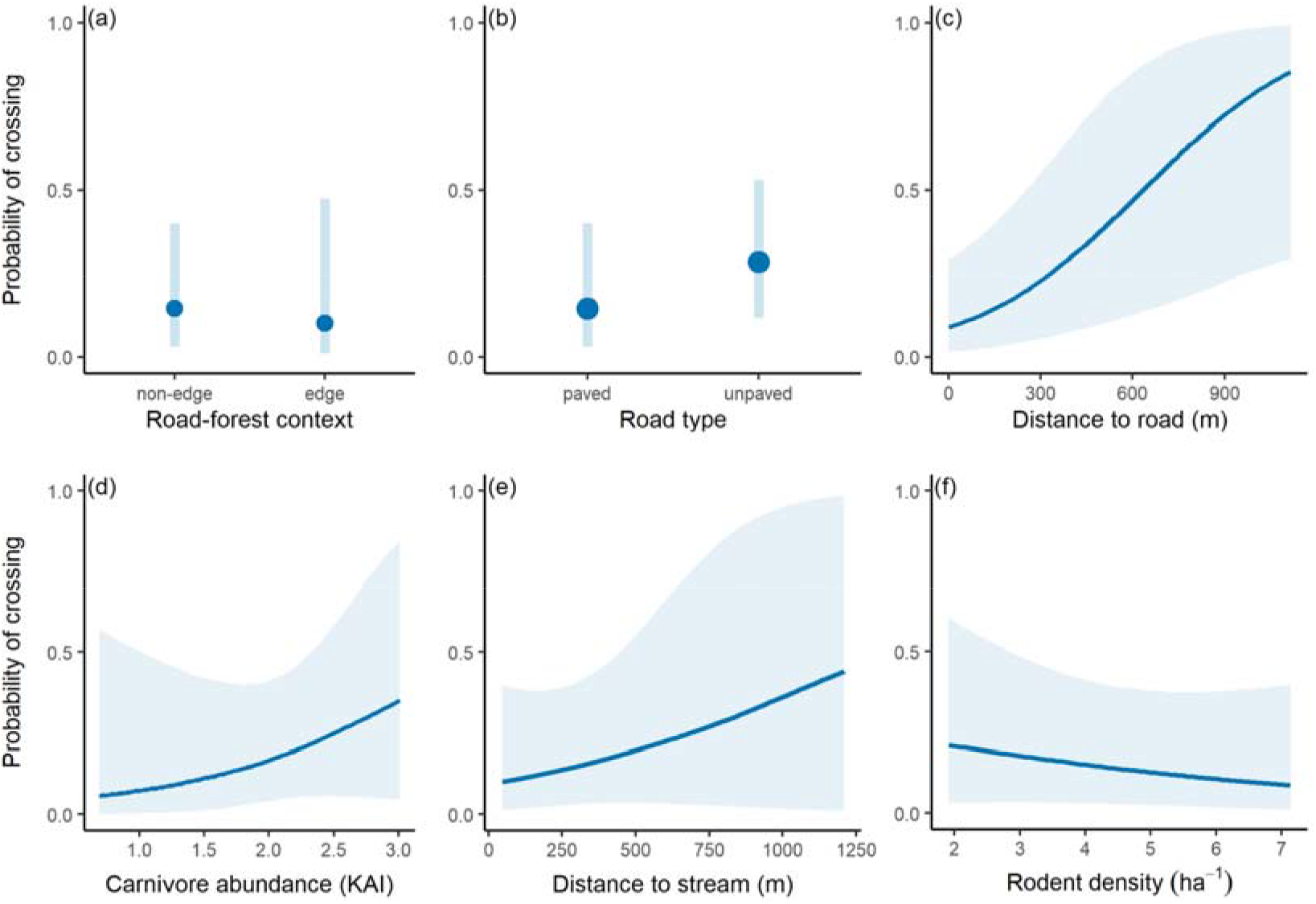
Mean fitted seed crossing probability (±95 % credible intervals) by road type (a), road-forest context (b) and covariates (c–f), as predicted by a Bayesian mixed-effects logistic model. See Table 1 for variable descriptions.

**Table 3.**
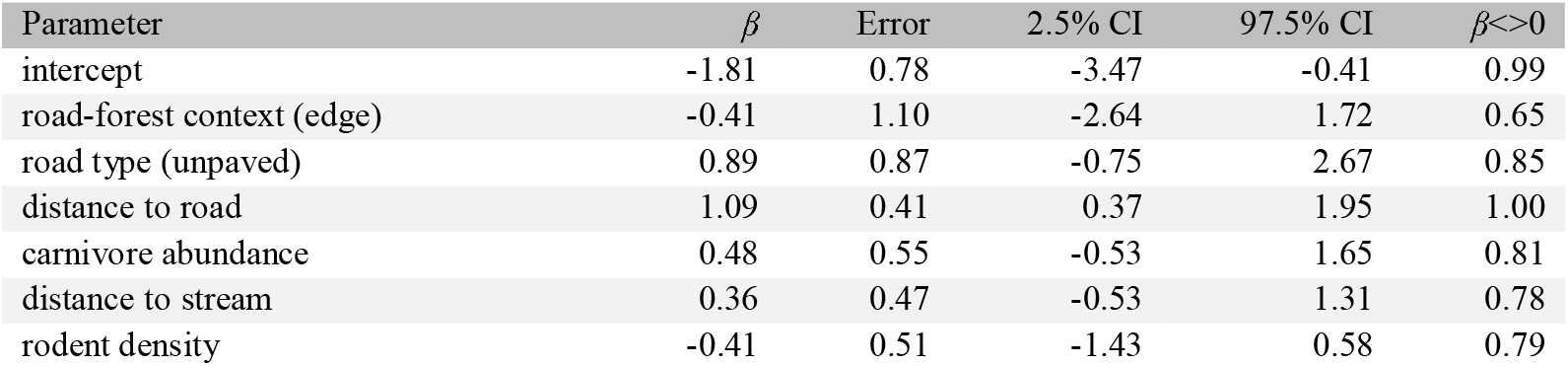
Summary of the fixed effects from the Bayesian mixed-effects logistic model evaluating the effects of road-forest context (non-edge vs. edge) and road-type (unpaved vs. paved) on carnivore-mediated seed crossing probability. Covariates include distance to road, carnivore abundance (carnivore kilometric abundance index; scats km□¹), distance to stream, and rodent density). CI = 95% credible interval; β<>0 = posterior probability that the effect is greater (or smaller) than zero, depending on the sign of the estimate. The potential scale reduction factor (Rhat) was 1.00 for all parameters.

**Table 4.** Fixed part of the Bayesian logistic model predicting the effect of hotspot presence on the probability of seed crossing mediated by carnivores. CI = 95% credible interval; β<>0 = posterior probability that the effect is greater (or smaller) than zero, depending on the sign of the estimate. The potential scale reduction factor (Rhat) was 1.00 for all parameters.

| Parameter | $\beta$ | Error | 2.5% CI | 97.5% CI | $\beta < > 0$ |
| --- | --- | --- | --- | --- | --- |
| intercept | -1.59 | 0.82 | -3.36 | -0.16 | 0.99 |
| carnivore roadkill hotspot (yes) | 1.62 | 1.20 | -0.64 | 4.06 | 0.92 |

Among covariates, distance to the road showed the clearest signal, with essentially all posterior mass above zero. Illustratively, model predictions indicate that a seed deposit located 100 m from the road had 33% higher probability of being a crossing than one only 10 m from the road (Fig. 3c). Crossing probability also tended to increase with carnivore abundance (KAI) and with greater distance to the nearest stream, and to decrease with rodent density (directional posterior probabilities of 0.81, 0.78, and 0.79, respectively). For illustration, model-based predictions show a 133% relative increase in crossing probability when KAI rises from 1 to 2 scats km□¹, a 43% increase when stream distance rises from 250 to 500 m, and a 27% decrease when rodent density increases from 2 to 4 individuals ha□¹ (Fig. 3d–f).

### Carnivore roadkill hotspots versus seed-crossing sections

We detected seven carnivore roadkill hotspots along the network of surveyed roads (Fig. 4). Of the seven hotspots, three intersected three of the six paved 1.2-km seed-crossing sections (one per section), and four occurred elsewhere on the network. Accordingly, the six seed-crossing sections were classified as hotspot-present (n = 3) or hotspot-absent (n = 3).

**Fig. 4.**
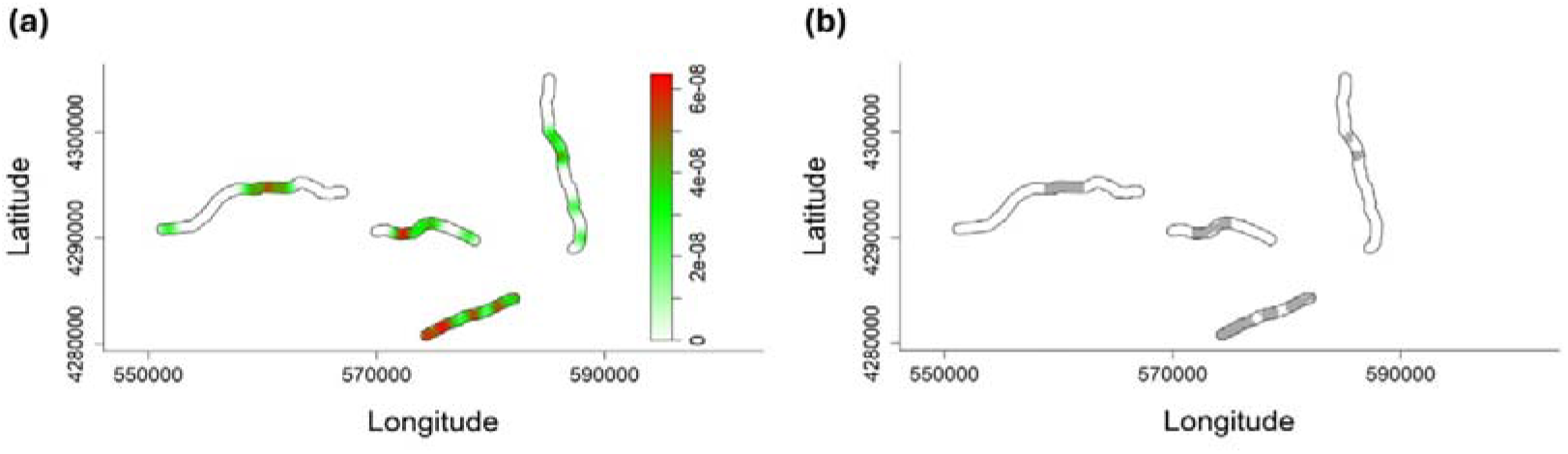
Detection of carnivore roadkill hotspots. (a) Kernel density estimation of roadkill intensity along the weekly-surveyed road network (warmer colours = higher density). (b) Statistically significant hotspots (95% Monte Carlo test against spatial randomness) shown as grey patches. Coordinates: EPSG:32629 (WGS 84 / UTM zone 29N).

Our Bayesian logistic regression on hotspot presence supported higher crossing probabilities on hotspot-present vs. hotspot-absent sections, with an estimated 92% directional posterior probability for this effect. Model predictions indicate a mean crossing probability of 0.51 on hotspot-present sections versus 0.18 on hotspot-absent sections—an absolute increase of 0.33 (+183% relative).

**Fig. 5.**
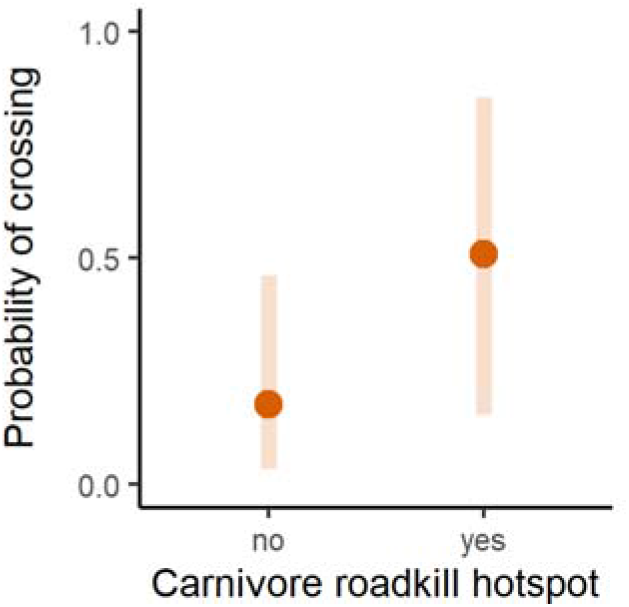
Mean fitted seed crossing probability (±95 % credible intervals) by carnivore roadkill hotspots, as predicted by a Bayesian logistic regression.

## DISCUSSION

Beyond viewing roads solely as barriers (Suárez-Esteban et al. 2013a; Ferreira et al. 2022), we show that carnivore-mediated seed crossings vary with road attributes—especially surface type—and provide the first evidence that sections with elevated seed-crossing probability coincide with carnivore roadkill hotspots. Together with previous evidence that roads alter animal-mediated seed dispersal and other interspecific interactions (Suárez-Esteban et al. 2013a; Chen et al. 2019; Quiles and Barrientos 2024; Craveiro et al. 2025), our findings indicate that road networks can modulate plant–animal interactions, with potential implications for connectivity, restoration, and road planning.

Raw seed-crossing proportions were higher in non-edge than edge sections, yet Bayesian inference offered only weak evidence for a road–forest context effect, suggesting any context signal is modest relative to road type. Still, habitat structure guides carnivore movement and crossing locations (Fabrizio et al. 2019; Valerio et al. 2019; Frangini et al. 2022): carnivores can be more active where continuous forest provides cover and resources on both sides (Tigas et al. 2002; Craveiro et al. 2025) and avoid open, altered habitats (Grünewald et al. 2010; Costa et al. 2022). For instance, red foxes—30% of our dispersal events—though generalists, often use forest more than open habitats (Cagnacci et al. 2004; Van Etten et al. 2007), consistent with the forest–open contrast at our edge sites. Pasture and cropland bordering edge roads likely diminish carnivore activity and thus reduce crossings, although generalists may also exploit edges (Andersen et al. 2017).

Road surface played the larger role. Despite roads being less of a barrier to carnivores than to many vertebrates (Craveiro et al. 2025), paved roads had fewer crossings than unpaved roads—plausibly due to heavier traffic and avoidance—whereas lower disturbance on unpaved roads may facilitate movements that promote seed dispersal (Suárez-Esteban et al. 2013b; Bueno et al. 2021).

Beyond road type, additional factors also shaped seed crossings. The positive distance-to-road effect is consistent with long gut-retention coupled with cumulative movement: seeds ingested near roads are more likely to be carried across as animals continue travelling (Grilo et al. 2015). Crossing probability also increased with carnivore activity (KAI), matching an encounter-rate expectation—more carnivores yield more dispersal events and thus more opportunities to cross—and plausibly reflecting social and spatial processes (territorial patrols, avoidance of dominants, niche partitioning) that extend movements (Doncaster and Macdonald 1991). Resource concentration along verges (e.g., carcasses, small mammals) can further intensify itinerancy and encounters within the carnivore guild (Galantinho et al. 2020; Craveiro et al. 2025). Although KAI is a scat-based activity index rather than a direct abundance measure and is sensitive to detectability, it nonetheless supports a plausible behavioural pathway from higher carnivore activity to higher seed-crossing probability.

Distance to streams and rodent density were additional but weaker effects (positive and negative, respectively). Streams can channel mesocarnivore movement by providing guidance, cover and resources (Grilo et al. 2008, 2016; Delgado et al. 2018; Craveiro et al. 2019; Azedo et al. 2022), keeping animals on one roadside and lowering crossing odds; as stream distance increases, movements are less constrained by riparian corridors and more likely to cut across roads, raising seed-crossing probability. This mechanism likely depends on riparian condition—degraded strips can disrupt movement and weaken seed dispersal along streams (Grilo et al. 2011). In turn, higher rodent densities near verges may reduce the incentive to risk crossing, effectively anchoring foraging to a single roadside (Ruiz-Capillas et al. 2013; Galantinho et al. 2020). This is consistent with carnivores co-occurring in prey-rich habitats and ranging more widely where prey is scarce (Šálek et al. 2014), and with trophic partitioning whereby stone martens shift diet under low small-mammal abundance to avoid competition with genets (Barrientos and Virgós 2006). Together these processes can depress cross-road movement when prey is abundant near verges—and promote it when prey is patchy—with cascading consequences for plant regeneration via predator–prey–seed links (Roemer et al. 2009).

Hotspots concentrated risk where seed crossings were frequent. Notably, two of the three hotspot-intersecting seed-crossing sections were non-edge (with forest on both sides of the road), whereas two of the three non-hotspot sections were edge. This pattern is consistent with continuous forest promoting both movement and mortality, pointing to paved, non-edge segments as priority targets for mitigation. Resource availability and connectivity likely draw carnivores to such places (Fabrizio et al. 2019; Valerio et al. 2019; Frangini et al. 2022), potentially making them focal points for plant–animal mutualisms. Consistently, European polecat roadkills were more frequent where natural habitats bordered roads in Spain (Carmona et al. 2024), and in southern Portugal—partly overlapping our area—favourable habitat predicted carnivore roadkills with a reported rate of 47 roadkills per 100 km yr□¹ (Grilo et al. 2009), 28% below our estimate. Mitigation at hotspots is crucial to reduce mortality while retaining seed-dispersal functions. Notwithstanding, forest edges are also important, as these transition zones may facilitate seed movement into altered habitats, potentially contributing to habitat restoration.

Importantly, dispersal may continue after a disperser’s death: carcasses subsidize scavengers (Mata et al. 2017), which can ingest and move seeds from roadkilled animals (Hämäläinen et al. 2017). However, this post-mortem dispersal is complementary and cannot fully replace primary dispersal by the live animal; roadkill also removes the disperser from the system, with a time-extended cost in foregone future dispersal events.

Although we sampled across all four seasons to minimize seasonal bias, roadkill risk also varies with sex, age, and species (Grilo et al. 2009; Ascensão et al. 2014; Carvalho et al. 2018). For instance, on a motorway in southern Portugal, 69% of roadkilled genets were subadults, with mortality peaking during the dispersal season (Carvalho et al. 2018), whereas in our national-road data most carcasses were adults. Such age- and season-specific patterns warrant attention because they can disproportionately affect recruitment and the ecological roles of dispersers, with consequences for long-term resilience.

These findings should be interpreted in light of the spatial and ecological scope of our study. The experiment was conducted in Mediterranean oak woodlands, and the hotspot comparison involved six paved sections. The magnitude and generality of the observed relationships may therefore differ in landscapes with other habitat structures, carnivore communities, road characteristics, and traffic regimes. Replication across broader road networks and contrasting ecosystems is needed to assess the generality of the association between roadkill hotspots and seed crossings and the transferability of hotspot-based mitigation.

### Conservation implications

Our results point to a clear priority: treat first the paved road segments that overlap carnivore roadkill hotspots. These are the places where risk and ecological function co-occur—seed crossings are most likely and mortality is concentrated. Targeting these segments yields dual benefits: fewer deaths and maintenance of carnivore-mediated seed flow, a key pathway of functional connectivity in many managed landscapes (Genes and Dirzo 2022).

Mitigation at these sites should be hotspot-guided, not diffuse. In practice, combine: (i) speed management tailored to short sections (e.g., dynamic signage, targeted enforcement; Jägerbrand et al. 2018); (ii) directional fencing that funnels animals to existing structures—culverts retrofitted with dry ledges/benches (Craveiro et al. 2019) and, where warranted, overpasses (Azedo et al. 2022); and (iii) verge management to reduce attractants (prompt carcass removal; vegetation and waste practices that limit small-mammal peaks) (Planillo et al. 2018) while retaining essential cover. Where culverts already exist, modest retrofits plus smart fencing are cost-effective and rapidly deployable (Craveiro et al. 2019).

Landscape context matters. Riparian strips can channel movement; where streams meet roads, pairing hydraulic works with fauna passages improves crossing safety without sacrificing guidance/cover (Grilo et al. 2008). Where rodents are abundant near verges, foraging may be fixed to one roadside, reducing crossings but potentially raising exposure to vehicles—another reason to align vegetation and waste management with mitigation goals. A light-weight monitor–act–reassess workflow—kernel-density roadkill mapping + seed-mimic trials + cameras—offers a replicable protocol to detect hotspots, implement measures, and evaluate effectiveness (ideally in BACI designs). Update hotspot maps regularly and after interventions.

The hotspot-first approach is transferable to other roaded mosaics where mesocarnivores disperse seeds. While evidence was strongest for road surface and more limited for road–forest context and other covariates, the expected gains and co-benefits justify pilots on paved, non-edge segments with confirmed hotspots, coupled with adaptive monitoring. This aligns road-safety and conservation mandates and offers a pragmatic route to reconcile mobility infrastructure with living connectivity.

