## Supporting Information for "Carnivore-mediated seed crossings are more likely on roadkill hotspots"

**Appendix A**

**Table A.1** Counts of carnivore roadkills by species within KDE-defined hotspots intersecting the paved 1.2-km seed-crossing sections, and elsewhere along the roadkill transects. “Seed-crossing sections” refer only to paved sections intersecting a hotspot (no roadkills occurred in seed-crossing sections without hotspots).

| Species | Seed-crossing sections | Elsewhere | *Total* |
| --- | --- | --- | --- |
| *Vulpes vulpes* | 2 | 9 | 11 |
| *Meles meles* | 6 | 1 | 7 |
| *Martes foina* | 2 | 3 | 5 |
| *Herpestes ichneumon* | 2 | 3 | 5 |
| *Genetta genetta* | 3 | 1 | 4 |
| *Lutra lutra* | 0 | 1 | 1 |
| *Total* | 15 | 18 | 33 |
